# Elevated Delta Power in a Maternal *UBE3A*-Deletion Pig Model of Angelman Syndrome

**DOI:** 10.64898/2026.02.23.707227

**Authors:** Luke S. Myers, Tanner G. Monk, Luis A. Martinez, Alasdair J. Taylor, Sarah G. Christian, Anne E. Anderson, Scott V. Dindot

## Abstract

Angelman syndrome is a neurodevelopmental disorder caused by loss of the maternally inherited *UBE3A* allele and is characterized by severe cognitive, motor, and communication impairments. Increased delta (1– 4 Hz) activity on electroencephalogram (EEG) assessed by visual inspection and by spectral power analysis is a robust feature of the disorder in humans and rodent models and is used as a biomarker of Angelman syndrome. This aspect of the phenotype has not been evaluated in the recently developed pig model of Angelman syndrome. Here, we analyzed scalp EEG recordings from freely moving pigs carrying a maternal *UBE3A* deletion (*UBE3A*^-/+^) across three age groups to determine whether they recapitulate the delta power abnormalities characteristic of the disorder. *UBE3A*^-/+^ pigs exhibited elevated delta power during both wakefulness and sleep compared with wild-type littermates, with the largest differences observed during the awake state. The typical increase in delta power that accompanies the transition from wakefulness to sleep was also reduced in *UBE3A*^-/+^ pigs. These effects were observed across study groups, demonstrating that the maternal *UBE3A*-deletion pig model reproduces the elevated delta power EEG phenotype of Angelman syndrome. Our results establish noninvasive scalp EEG as a translationally relevant tool for assessing neural dysfunction in this large-animal model and provide a framework for preclinical therapeutic testing. This work strengthens the utility of the pig model for mechanistic studies and therapeutic development in Angelman syndrome.

## Introduction

Angelman syndrome is a rare neurodevelopmental disorder affecting 1 in 10,000–24,000 live births^(1)^. Individuals present with intellectual disability, severe speech impairment, ataxia, motor incoordination, epilepsy, and microcephaly^(2)^. The most common etiology is a maternal deletion of the 15q11–q13 region that encompasses *UBE3A*^(3)^. In central nervous system (CNS) neurons, only the maternal *UBE3A* allele is expressed, so maternal deletion eliminates neuronal UBE3A and causes Angelman syndrome^(3)^.

A large-animal model of Angelman syndrome was developed using CRISPR–Cas9 to generate pigs (*Sus scrofa*) with a maternal *UBE3A* deletion^(4)^. This model enables studies in an organism anatomically closer to humans and with a more comparable neurodevelopmental trajectory than traditional rodent models^(5-9)^. Numerous hallmark Angelman syndrome phenotypes have been reported in these pigs, including hypotonia, ataxia, altered and reduced vocalizations, feeding difficulties, and reduced brain size^(4)^.

One robust Angelman syndrome biomarker used to index therapeutic response in both patients and rodent models is increased delta power on electroencephalography (EEG); however, this phenotype has not yet been characterized in the pig model. EEG records electrical activity of the cerebral cortex, providing insight into brain neurophysiology. In humans, over 80% of individuals with Angelman syndrome show increased delta (1–4 Hz) activity on EEG^(2, 10, 11)^. Rodent models of Angelman syndrome also exhibit elevated delta activity^(10-13)^. In both individuals with Angelman syndrome and rodent models, delta spectral power is elevated at younger ages, declines with maturation, yet remains persistently higher than controls throughout life^(10, 12)^. Therefore, assessing multiple preclinically relevant ages in the pig model is necessary to capture the developmental trajectory of delta power. These findings support the use of EEG delta metrics as quantitative biomarkers and outcome measures for therapeutic development in the pig model.

In humans with Angelman syndrome, EEG assessments are commonly performed with scalp electrodes^(14)^. In rodent models, EEG assessments are typically invasive and require craniotomies for intracranial electrode implantation because the skull is too small for scalp systems^(10)^. Pigs, however, share closer cranial anatomy and size with humans, permitting the use of scalp electrodes^(15)^. This enables noninvasive, repeatable recordings in freely moving awake pigs and avoids implantation- or surgery-related confounds. By aligning electrode type and placement with clinical practice, the pig model improves comparability and strengthens the translational value of EEG findings.

Here, we recorded scalp EEG from freely moving wild-type pigs and pigs with a maternal *UBE3A* deletion across three age groups. We observed increased relative delta power during wakefulness and sleep in pigs with a maternal *UBE3A* deletion compared with wild-type controls and outlined practical challenges specific to scalp EEG in freely moving pigs. These results support the potential of scalp EEG as an informative tool for preclinical Angelman syndrome research in a large-animal model.

## Methods

### Animal model and housing conditions

All experimental procedures were approved by the Institutional Animal Care and Use Committee (IACUC) at Texas A&M University. Yorkshire–Landrace pigs carrying a maternal *UBE3A* deletion (*UBE3A*^-/+^) and their wild-type littermates (WT) were used.

Pigs were housed in climate-controlled rooms maintained at 74 ± 1°F, with a 12-hour light/dark cycle and 40–60% relative humidity. Until 40 days of age, feed was provided ad libitum; thereafter, pigs received 4% of their body weight daily. Animals were group-housed, except when temporary isolation was required for health reasons. To ensure consistency in recordings, all EEG sessions began between 8:00 AM and 10:00 AM (with lights turning on at 6:00 AM) and concluded by 1:00 PM.

No pigs received routine or consistent medical or pharmacological treatment for Angelman syndrome phenotypes. Interventions were administered only when necessary, such as under veterinary guidance or in response to life-threatening conditions. All animals were confirmed to be in good health prior to EEG recordings.

### Sample size and exclusions

Enrollment was dictated by breeding yield and funding availability. A total of 14 pigs were included in the study (WT n = 6, 3F/3M; *UBE3A*^-/+^ n = 8, 4F/4M). To reflect developmental differences, animals were stratified into predefined age bins: 4–7 weeks (P30–P48), 9–11 weeks (P60–P76), and 13–17 weeks (P90– P120). The final sample sizes contributing usable EEG data at each age group were: 4–7 weeks: WT n = 6 (3F/3M), *UBE3A*^-/+^ n = 7 (4F/3M); 9–11 weeks: WT n = 4 (1F/3M), *UBE3A*^-/+^ n = 4 (1F/3M); 13–17 weeks: WT n = 4 (1F/3M), *UBE3A*^-/+^ n = 5 (2F/3M).

Exclusion criteria were prespecified and applied uniformly. Recording sessions were terminated if a pig removed electrodes ≥3 times within 5 minutes, more than 5 times during the session, or if the transmitter or leads were damaged. Depending on the severity of the disruption or equipment damage, animals were excluded from future sessions to protect animal welfare and prevent further equipment loss. Sessions were also excluded if synchronized video recordings were unavailable. All termination and exclusion decisions were made before genotype unblinding.

### Genotyping

Genomic DNA was isolated from tissue using the DNeasy Blood & Tissue Kit (Qiagen, 69504) with overnight lysis at 56 °C on a rocker. DNA was eluted in 200 µL AE buffer and quantified on a Qubit 4 fluorometer using the dsDNA BR Assay (Thermo Fisher Scientific, Q32850). PCR genotyping reactions (20 µL) contained 25–50 ng DNA, 500 nM each of primers UBE3A 1-13F and UBE3A 1-13R2, and 10 µL 2× Phire Mix (Thermo Fisher Scientific, F126L). Amplification was performed on a Bio-Rad T100 thermal cycler with the following program: 98 °C for 30 s; 35 cycles of 98 °C for 5 s, 64 °C for 5 s, and 72 °C for 5 s; and a final extension at 72 °C for 1 min. PCR products were analyzed by agarose gel (1.5% agarose in 1× TBE) electrophoresis. Each run included a no-template control and positive controls from known WT and *UBE3A*-deletion carriers.

### EEG acquisition hardware

EEG was recorded with a non-invasive eegPACK / easyTEL+ RP wireless telemetry system (emka TECHNOLOGIES) configured by manufacturer for swine. The system used a wireless eegPACK transmitter powered by two zinc-air PR13 (1.45 V) cells installed at the start of each session; no battery swaps occurred mid-session. The transmitter acquires up to four biopotential channels with user-selectable sampling to 500 Hz and input ranges of ±2 mV. Per manufacturer specifications, the biopotential front end is AC-coupled with hardware bandwidth of approximately 1–150 Hz.

Signals were carried from the scalp electrodes to the transmitter via 0.3048 m (1 ft) stainless-steel, polyamide-insulated leads that were color-coded by input. Leads terminated in snap-type, monopolar disposable surface electrodes (Pediatric Foam Gel ECG 38 mm MDSM611050H, Medline).

A single emka digitalRECEIVER, positioned ≤3 m from the recording pen, simultaneously received data from 1–4 transmitters and relayed it to the acquisition workstation over a 20 m RJ45 (Cat-5e U/UTP) ethernet run. Data were captured on a Dell workstation (Intel vPro i7).

### EEG acquisition program and settings

EEG was acquired with an emka TECHNOLOGIES wireless eegPACK transmitter controlled by IOX software (version 2.10.38; April 11, 2022). The digital telemetry configuration was v1.2.1.48 (January 7, 2022). In our recordings, the sampling rate was 500 Hz, the input range was ±2,000 µV, and ADC resolution was 10-bit (≈3.9 µV/LSB). Baseline removal was disabled, and no online high-, low-, or notch filters were applied during acquisition. Following data collection, the native IOX files (.iox) were exported as European Data Format (EDF) for analysis in LabChart 8.

### Video acquisition and synchronization

Each pen was recorded with a GoPro HERO7 camera mounted in the top corner to provide a downward view of the pig. Video recording ran continuously during EEG sessions at 1440p (60 fps). Manual synchronization to EEG was performed using audible cues recorded on the video at session start (and as needed), and alignment was applied during analysis.

### EEG electrode placement

Electrode placement was performed under isoflurane anesthesia to ensure animal comfort and precise positioning. Isoflurane was delivered via face mask in oxygen at 1 L/min (induction 5%; maintenance 2– 3%). Anesthetic exposure typically lasted 6–12 minutes. Pigs were allowed a 60-minute washout. No epochs were taken or analyses performed during these washout periods. Pigs aged 4–7 weeks were held during isoflurane induction and then transferred to a surgical table for electrode attachment, whereas pigs aged 9–11 and 13–17 weeks were placed in a restraint sling for the entire procedure.

To optimize electrode adhesion, minimize signal interference, and ensure consistent contact, the scalp was shaved to remove all hair. The skin was sequentially cleaned: first with a warm, damp gauze to remove debris, followed by alcohol wipes to eliminate oils, and finally with a dry gauze to remove any residual moisture or alcohol. After cleaning, anatomical landmarks were used for standardized electrode placement. Lines were drawn from the tear ducts to the base of the adjacent ear, and their intersection served as a reference point approximating the frontal cortex.

Pigs were fitted with four biopotential electrodes positioned at the anterior left, anterior right, posterior left, and posterior right, with a midline ground electrode placed posteriorly. EEG signals were acquired in a referential (monopolar) montage, with each active electrode recorded relative to a common system reference. The midline electrode served solely as the ground connection to stabilize the amplifier and reduce common-mode noise; loss of this electrode disrupted all channels, but it was not used as a signal reference. For the present study, analyses focused on the anterior left and posterior right channels. Electrode adhesive was trimmed as needed to accommodate scalp size while maintaining a 2–3 mm margin around the conductive gel to preserve signal integrity.

Following electrode placement, the transmitter and cables were secured using self-adhesive bandage wrap. The transmitter was positioned between the shoulder blades, and the bandage wrap was carefully applied to anchor the cables along the back. To minimize movement-related artifacts, no material was placed directly over the electrodes. Once all equipment was properly secured, isoflurane administration was discontinued, and the pig was transferred to a pen for EEG recordings.

### EEG recording conditions

During EEG recordings, each pig was placed alone in a clean pen designed to resemble its home pen, with identical temperature and lighting conditions. Multiple EEG recordings were conducted simultaneously, with two to six pigs housed in the same room. Each pig was individually housed in a manner that prevented visual contact with others.

Recordings were intended to last three hours (±30 minutes). However, in some cases, recordings were shortened due to pigs learning to continuously remove their electrodes or damaging equipment. Throughout the recording period, lights remained on, and personnel were not present in the room except when necessary to reattach electrodes.

### EEG epoch selection

All EEG reviews and analysis were performed with investigators blinded to genotype. Traces were reviewed in LabChart 8 (ADInstruments, v8.1.30) using both the time-series and spectrogram views. Recordings contained frequent movement- and muscle-related artifacts, so clean epochs were identified by visual scanning the time series for artifact-free segments. Candidate epochs ranged from 30 s to 5 min and were required to be free of line interference, cable movement, chewing or muscle bursts, abrupt high-amplitude transients, amplifier saturation, or sustained broadband elevations. Screening began on the posterior right electrode, the channel least affected by facial musculature, and the same time intervals were then inspected on the anterior left electrode to verify that both channels were clean. Epochs were retained only when channels showed a stable baseline and clean spectrographic structure.

### Behavioral state classification

Behavioral state for each retained epoch was classified from synchronized video by investigators blinded to genotype using eye behavior, posture, and movement. Spectrogram visualization was used as an adjunct to confirm that the selected epochs were physiologically consistent with the observed behavioral state (e.g., sustained low-frequency dominance during sleep) but was not used as a primary classification criterion. Four states were initially defined: lying-alert, lying-not alert, standing, and moving. Standing and moving produced excessive motion artifacts, yielding too few clean epochs for analysis. Therefore, all analyses focused on the two lying states, corresponding to the awake (lying—alert) and asleep (lying—not alert) vigilance states used in the Results.

### Spectral analysis

Spectral analysis was performed in LabChart 8 (v8.1.30, 7/25/2024) using the Power Spectrum module. Power spectral density was computed using Welch’s method (Hann window; 0.512-s segments [256 samples at 500 Hz]; 50% overlap). Spectra were computed using a 512-point FFT (zero-padded), yielding a frequency bin spacing of 0.9766 Hz and a single-sided spectrum. For each accepted epoch and channel, integrated broadband power (0–50 Hz) was calculated as the sum across frequency bins ≤50 Hz. Relative delta power was defined as the sum from 0.98–3.91 Hz (≈1–4 Hz) divided by the 0–50 Hz broadband power within the same epoch. A 60-Hz notch was not applied because the analysis band ended at 50 Hz. On rare occasions, narrowband interference at ∼20 or ∼40 Hz was present; a narrow notch centered at the interfering frequency was applied to those epochs only.

### Statistics

The sample size was determined based on available funding and logistical feasibility rather than a formal statistical power calculation. While this approach may limit the statistical power of our findings, we ensured that the number of animals used was sufficient to provide meaningful biological insights while adhering to ethical guidelines for animal research. All statistical analyses were performed in JMP Pro 18 (Student Edition), and statistical significance was defined as p < 0.05. Outliers were identified using the interquartile range (IQR) criterion (values >1.5 × IQR above the third quartile or below the first quartile) and were removed prior to analysis. Sex was not included as a biological variable due to the limited number of males and females per group. Genotype effects on relative delta power were assessed using linear mixed-effects models with genotype and age group as fixed factors and animal ID included as a random effect to account for repeated measures within animals. Percentage differences between genotypes were calculated using the WT mean as the baseline (100 × [*UBE3A*^-/+^ mean – WT mean] / WT mean). Percentage changes between vigilance states were calculated using the awake value as the baseline (100 × [asleep – awake] / awake).

## Results

A recently developed pig model of Angelman syndrome has been shown to recapitulate many of the disorder’s hallmark phenotypes, including several features not previously observed in other animal models^(4)^. However, whether this pig model also exhibits the characteristic increase in delta power on EEG as consistently reported in individuals with Angelman syndrome and in rodent models remains unknown^(12)^. To address this, we analyzed relative delta power in EEG recordings obtained from 14 Yorkshire–Landrace pigs (6 wild-type and 8 maternal *UBE3A* deletion [*UBE3A*^-/+^]) across three developmental age groups: 4– 7, 9–11, and 13–17 weeks old. These age ranges correspond to approximately 51–62%, 68–75%, and 81– 90% of maximal brain growth in pigs^(6)^, respectively. Genotypes were confirmed by PCR-based genotyping, verifying the presence of the maternal *UBE3A* deletion in *UBE3A*^-/+^ pigs.

Electrodes were positioned over the anterior left and posterior right scalp regions (**Figure 1A**), approximately corresponding to the frontal and occipital cortices, respectively, enabling assessment of regional differences in brain activity. EEG recordings were then obtained from freely moving pigs using a non-invasive wireless telemetry system configured for swine (**Figure 1B**). Following artifact rejection, a total of 346 individual epochs of good-quality EEG data (140 from wild-type and 206 from *UBE3A*^-/+^ pigs) were analyzed to evaluate the effects of a maternal *UBE3A* deletion on relative delta power in this pig model of Angelman syndrome.

**Figure 1.**
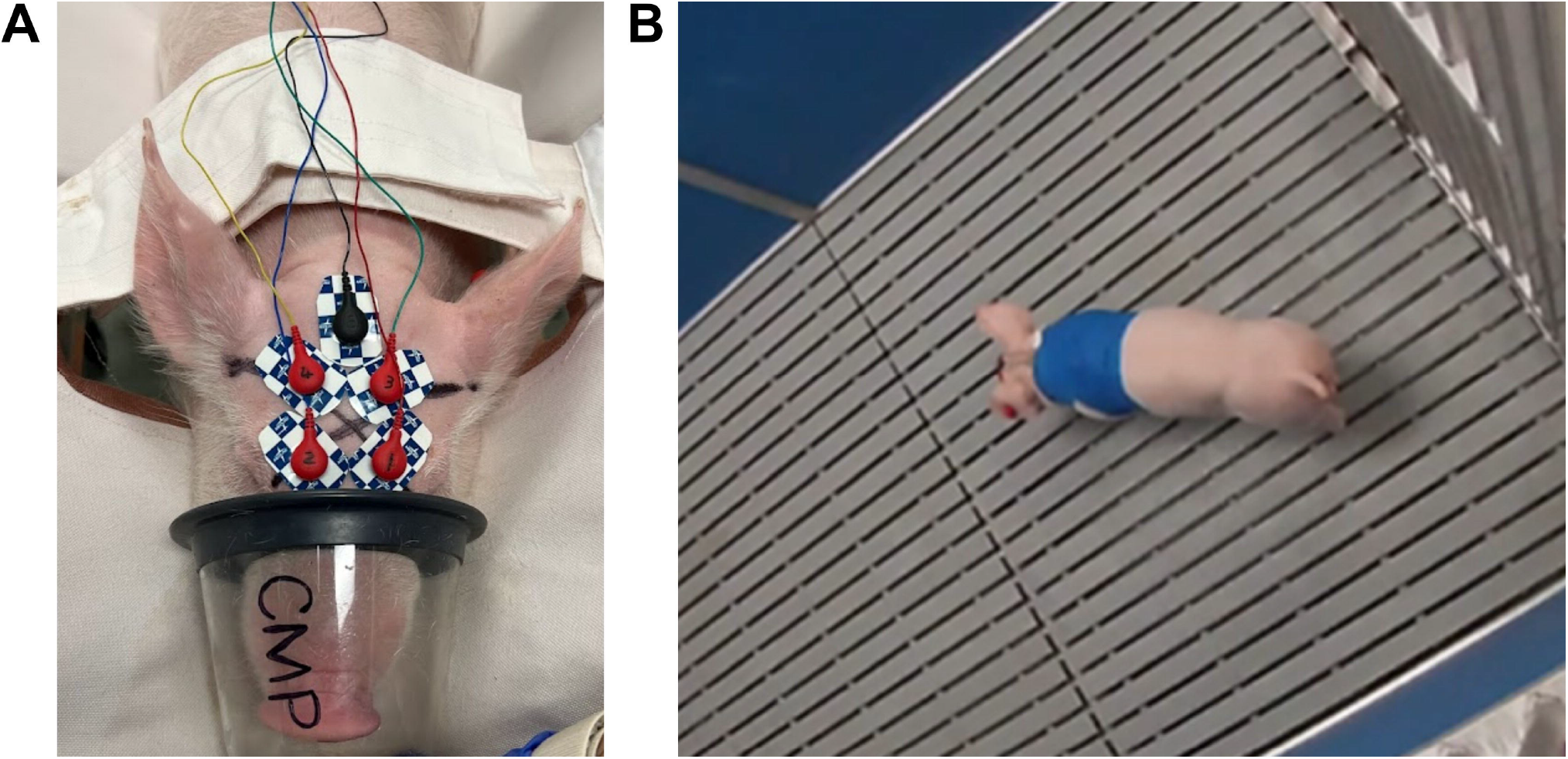
Representative images of noninvasive scalp EEG recordings in pigs. (**A**) Placement of adhesive scalp electrodes. Four electrodes were positioned on the scalp, but only the anterior left and posterior right channels were analyzed in the present study. (**B**) Pig undergoing a free-roaming EEG recording in a home-style pen, with the wireless transmitter secured between the shoulder blades using self-adhesive veterinary wrap and disposable adhesive scalp electrodes.

### Increased relative delta power in UBE3A^-/+^ pigs during the awake vigilance state

The greatest genotype-dependent differences in relative delta power were observed during the awake state (**Figure 2A-C**). Across all age groups, *UBE3A*^-/+^ pigs exhibited significantly higher relative delta power than wild-type pigs at the anterior left electrode (4–7 weeks: F_(1, 8.5)_ = 5.29, p = 0.0487; 9–11 weeks: F_(1, 7.8)_ = 37.90, p = 0.0003; 13–17 weeks: F_(1, 8.1)_ = 9.70, p = 0.0141; **Figure 2A**). At the posterior right electrode, a significant genotype effect was detected only in the 9–11-week group (F_(1, 5.3)_ = 27.02, p = 0.003), whereas the 4–7-week and 13–17-week groups showed nonsignificant trends in the same direction (**Figure 2B**).

**Figure 2.**
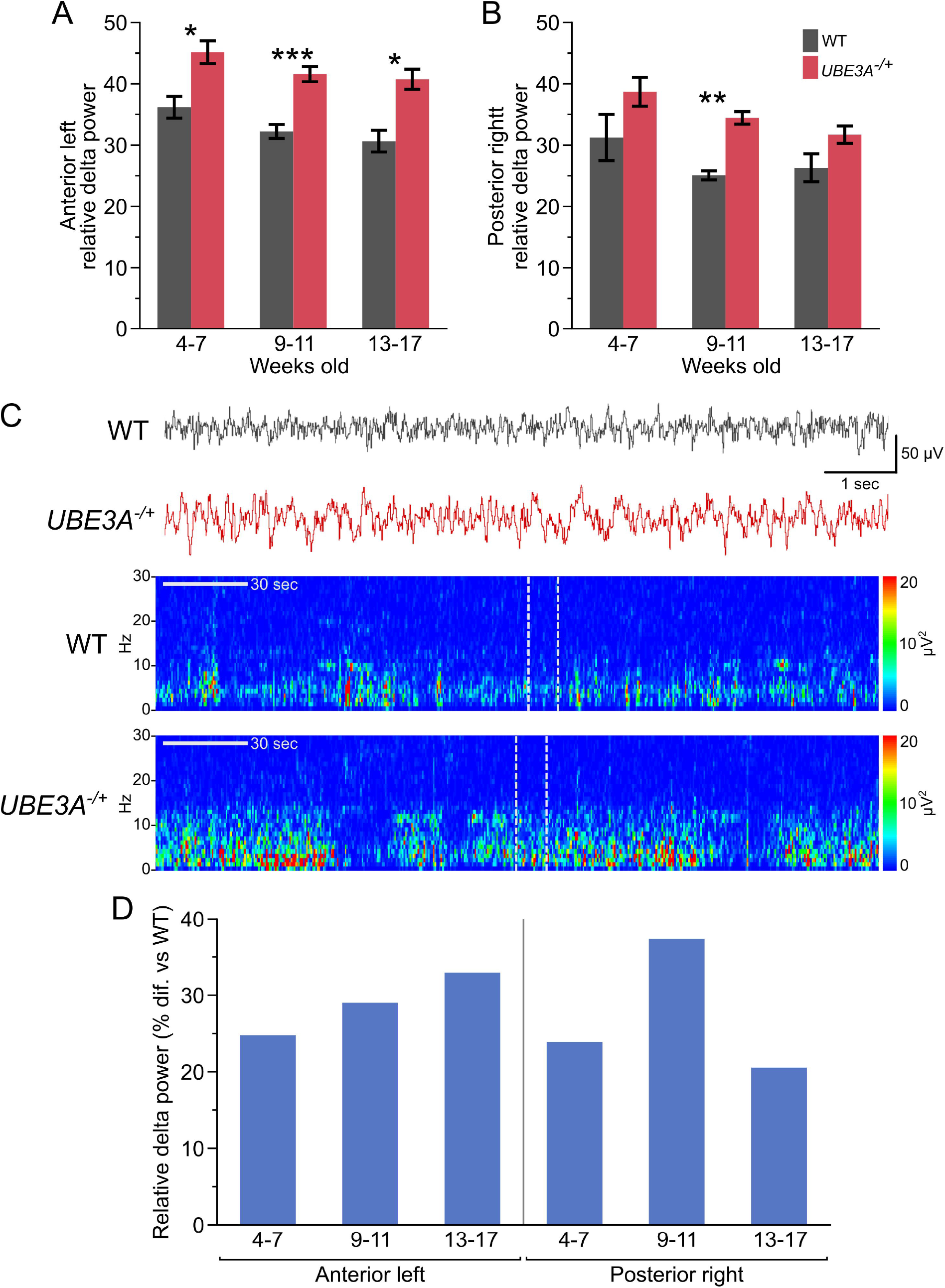
Increased relative delta power in *UBE3A*^-/+^ pigs during the awake state across development. (**A–B**) Relative delta power (1–4 Hz) at the anterior left (**A**) and posterior right (**B**) electrodes across three age groups (4–7, 9–11, and 13–17 weeks). Grey bars represent WT pigs and red bars represent *UBE3A*^-/+^ pigs. Data presented as mean ± SEM. (**C**) Representative EEG traces and corresponding spectrograms from 9–11-week-old WT (grey) and *UBE3A*^-/+^ (red) pigs. The EEG traces were taken from the epochs whose relative delta power was closest to each genotype’s group mean, ensuring representative visualization and minimizing selection bias. Dotted grey lines on the spectrograms indicate the segment from which the corresponding EEG trace was taken. (**D**) Percent difference in relative delta power between *UBE3A*^-/+^ pigs and age-matched WT controls during the awake state. Abbreviations: hertz (Hz), second (sec), volts (V), difference (Dif.), wild-type (WT) and maternal *UBE3A* deletion (*UBE3A*^-/+^). *P < 0.05, **P < 0.01, and ***P < 0.0001.

The largest percentage difference occurred at the posterior right electrode in pigs aged 9–11 weeks, where relative delta power was 37% higher in *UBE3A*^-/+^ pigs compared with wild-type controls (**Figure 2D**). The smallest genotype difference at this site was observed in the 13–17-week group, where *UBE3A*^-/+^ pigs still showed a 21% increase relative to wild-type pigs (**Figure 2D**). When averaged across ages, *UBE3A*^-/+^ pigs demonstrated an overall increase in relative delta power of 29% at the anterior left electrode and 27.3% at the posterior right electrode (**Figure 2D**).

### Increased relative delta power in UBE3A^-/+^ pigs during the asleep vigilance state

During the asleep state, *UBE3A*^-/+^ pigs also showed higher relative delta power than wild-type pigs at both electrode sites (**Figure 3A–C**), although the effect was less pronounced than during wakefulness. A significant genotype effect was detected only at the anterior left electrode in the 9–11-week group (F_(1, 20)_ = 10.47, p = 0.0041). Across all other ages and at the posterior right electrode, *UBE3A*^-/+^ pigs consistently exhibited higher delta power than wild-type pigs, but these differences did not reach statistical significance (**Figure 3A–B**).

**Figure 3.**
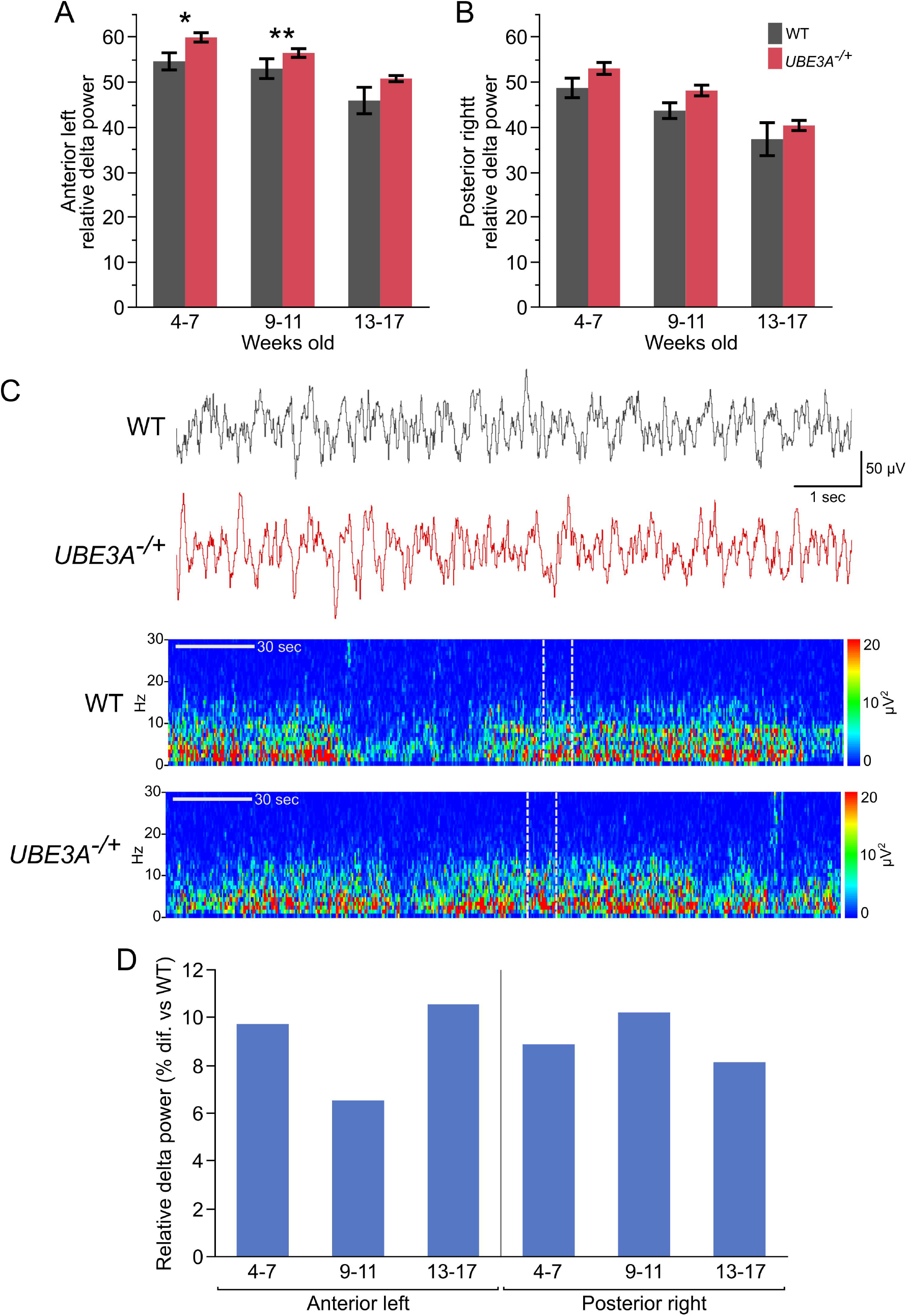
Increased relative delta power in *UBE3A*^-/+^ pigs during the asleep state across development. (**A–B**) Relative delta power (1–4 Hz) at the anterior left (**A**) and posterior right (**B**) electrodes across three age groups (4–7, 9–11, and 13–17 weeks). Grey bars represent WT pigs and red bars represent *UBE3A*^-/+^ pigs. Data presented as mean ± SEM. (**C**) Representative EEG traces and corresponding spectrograms from 9–11-week-old WT (grey) and *UBE3A*^-/+^ (red) pigs. The EEG traces were taken from the epochs whose relative delta power was closest to each genotype’s group mean, ensuring representative visualization and minimizing selection bias. Dotted grey lines on the spectrograms indicate the segment from which the corresponding EEG trace was taken. (**D**) Percent difference in relative delta power between *UBE3A*^-/+^ pigs and age-matched WT controls during the asleep state. Abbreviations: hertz (Hz), second (sec), volts (V), difference (Dif.), wild-type (WT) and maternal *UBE3A* deletion (*UBE3A*^-/+^). *P < 0.05, and **P < 0.01.

Despite limited statistical significance, *UBE3A*^-/+^ pigs showed modestly higher delta power across development. The largest percentage difference occurred at the anterior left electrode in the 13–17-week group, where *UBE3A*^-/+^ pigs showed an 11% increase relative to wild-type controls (**Figure 3D**). The smallest difference at this site was observed in the 9–11-week group, with a 7% increase (**Figure 3D**). Averaged across ages, *UBE3A*^-/+^ pigs demonstrated a 9.3% increase at the anterior left electrode and a 9.0% increase at the posterior right electrode compared with wild-type pigs (**Figure 3D**).

### UBE3A^-/+^ pigs exhibit a diminished delta power transition between awake and asleep states

When transitioning from the awake to the asleep vigilance state, wild-type pigs showed a markedly greater increase in relative delta power than *UBE3A*^-/+^ pigs (**Figure 4A-B**). In wild-type pigs, the largest increase was observed at the posterior right electrode in the 9–11 weeks age group, with relative delta power rising by 76% (**Figure 4B**). The same electrode and age group also exhibited the greatest increase in *UBE3A*^-/+^ pigs, but the change was smaller, at 40% (**Figure 4B**). Notably, this value was lower than even the smallest awake-to-asleep increase observed in wild-type pigs (42% at the posterior right electrode in the 13-17 weeks group). The smallest increase in *UBE3A*^-/+^ pigs occurred at the anterior left electrode at 13-17 weeks, with a 25% increase (**Figure 4A**). Across all age groups and electrodes, wild-type pigs exhibited an average 56.3% increase in relative delta power when transitioning from the awake to the asleep state, compared with an average increase of 33% in *UBE3A*^-/+^ pigs.

**Figure 4.**
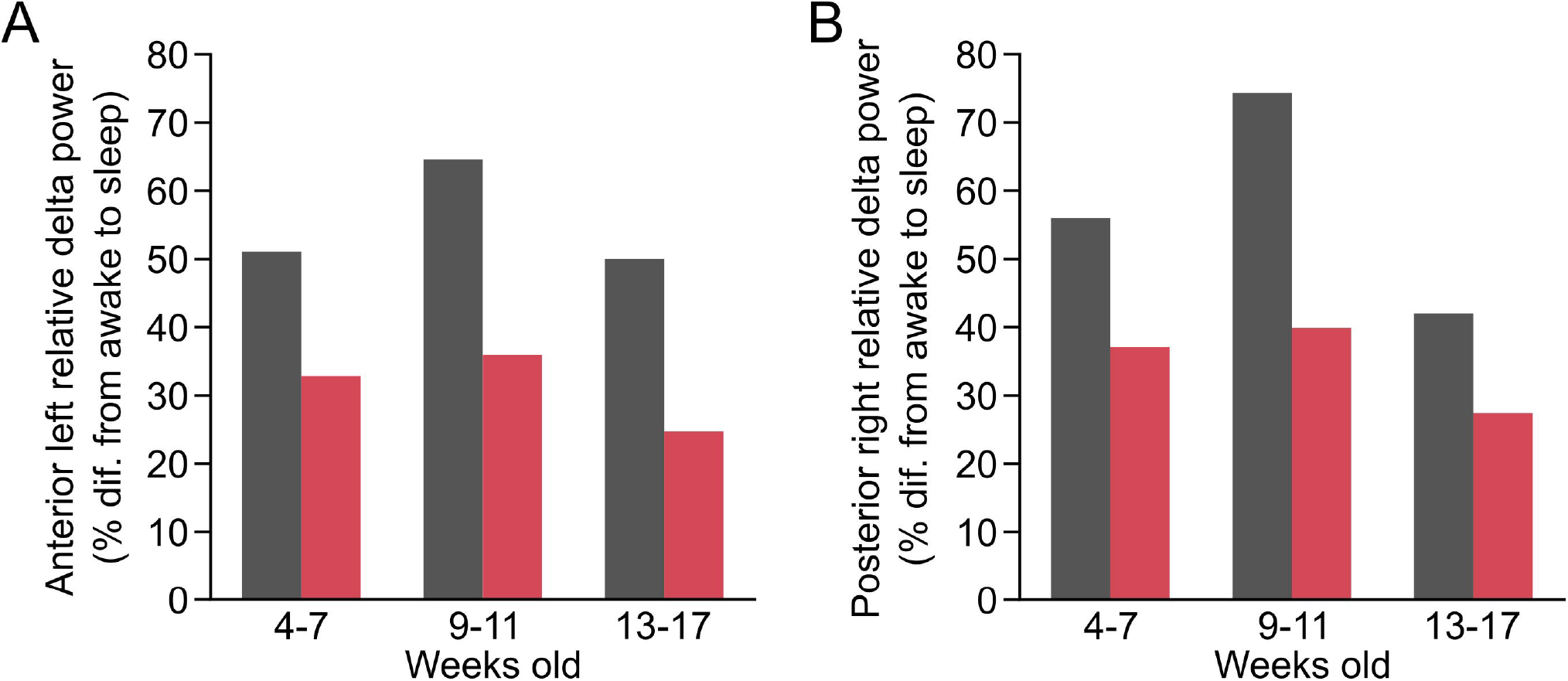
Reduced state-dependent increase in relative delta power in *UBE3A*^-/+^ pigs during the transition from awake to asleep. (**A–B**) Average percentage change in relative delta power when transitioning from the awake to the asleep state at the anterior left (**A**) and posterior right (**B**) electrodes across the three age groups. Grey bars represent WT pigs and red bars represent *UBE3A*^-/+^ pigs. Abbreviations: difference (Dif.), wild-type (WT) and maternal *UBE3A* deletion (*UBE3A*^-/+^).

Together, these findings demonstrate that pigs with a maternal *UBE3A* deletion consistently exhibit elevated relative delta power across vigilance states, with the greatest differences observed during wakefulness. Moreover, the typical increase in delta power from the awake to asleep state was attenuated in *UBE3A*^-/+^ pigs compared with wild-type controls. This blunted transition may reflect a ceiling effect due to already elevated baseline delta activity during wakefulness or impaired sleep–wake state regulation, a feature commonly reported in individuals with Angelman syndrome^(16, 17)^. Overall, these results confirm that the Angelman syndrome pig model reproduces the increased delta power characteristic of the disorder.

## Discussion

Abnormal increases in delta power are a well-established hallmark of Angelman syndrome, consistently reported across human studies and rodent models^(2, 10-13)^. In the present study, we extend these findings to a large-animal model showing that pigs with a maternal *UBE3A* deletion exhibit elevated relative delta power across vigilance states, with the largest differences observed during wakefulness.

EEG assessments are currently being used and explored as quantitative outcome measures in clinical trials for Angelman syndrome^(12, 18, 19)^. The ability to capture comparable EEG readouts in the pig model provides an opportunity to align preclinical and clinical outcome measures, thereby strengthening translational predictability. Pigs share closer neuroanatomical and physiological similarities to humans than rodents^(5-9)^, and the use of noninvasive scalp electrodes, rather than surgically implanted intracranial electrodes, permits recordings that more closely align with methods used to measure delta power in patients. This nonterminal approach also enables repeated, longitudinal assessments of brain function and cortical EEG in the same animal, eliminating the need for invasive EEG monitoring or euthanasia-based analyses such as microscopy, RNA, or protein quantification to assess therapeutic effects. As a result, researchers can monitor how cerebral EEG activity evolves over time and evaluate the progression or reversal of disease-related phenotypes following intervention. Furthermore, positioning multiple electrodes across the scalp allows investigation of regional treatment effects, which can be compared to human cortical patterns to identify brain regions or behavioral phenotypes that will potentially be more responsive to therapeutic rescue.

A notable strength, but also a limitation, of this study was the use of noninvasive scalp electrodes in freely moving pigs. The large size of the pig’s head allows placement of multiple adhesive electrodes in a configuration relatively similar to that used in clinical EEG, offering an opportunity for direct comparison between preclinical and human data. This alignment enhances translational relevance, particularly for evaluating therapeutic rescue effects. However, unlike implanted electrodes, adhesive scalp electrodes rely solely on medical-grade pressure-sensitive adhesives, making them more susceptible to movement artifacts and electrode displacement. As a result, recordings could not be reliably obtained while pigs were actively walking or moving, as these activities produced substantial motion-related noise that obscured the underlying neural signal.

This method of electrode attachment also introduced behavioral challenges. As the pigs became familiar with the recording setup, they learned to rub their heads against the pen walls to remove the electrodes. To minimize this risk, recording sessions were limited to approximately three hours. Longer sessions increased the likelihood of electrode removal, which could compromise both data quality and animal welfare, and in some cases precluded further recordings from that animal. Consequently, overnight EEG recordings could not be performed, and the frequency of artifacts limited reliable estimation of absolute spectral power. Therefore, analyses focused on relative delta power, which normalizes broadband variability and is less sensitive to impedance fluctuations and movement-related amplitude shifts in freely moving animals. Attempts to record younger pigs (5, 6, and 15 days old) were made but ultimately unsuccessful, as separation from the sow caused agitation and constant movement, preventing acquisition of artifact-free data sufficient for reliable analysis.

Several strategies were explored to improve electrode retention. We tested acclimation procedures by fitting pigs with electrodes and mock transmitters several days prior to recordings, with the goal of reducing novelty-driven removal behavior. However, this approach had the opposite effect, accelerating their ability to dislodge the equipment. Similarly, attempts to use elastic “hats” or additional medical-grade tapes to secure the electrodes increased the incidence of artifacts, as normal muscle movements transmitted motion across all electrodes simultaneously. Given these outcomes, we opted for shorter recording durations and avoided keeping the equipment on animals outside of active recording periods to limit learning opportunities. Four electrodes were placed on the pigs; however, only two were analyzed for relative delta power due to time and resource constraints. Consequently, future studies could consider attaching only two electrodes positioned more centrally on the scalp, reducing the pig’s ability to remove them. Studies should also be performed in older animals to determine whether this difference in relative delta power persists and whether the age-dependent reduction in delta power observed in individuals with Angelman syndrome and rodent models is also present in pigs.

Overall, despite the practical challenges associated with scalp EEG recordings in freely moving pigs, this study demonstrates that a maternal *UBE3A* deletion alters EEG delta activity in a manner consistent with the defining EEG phenotype of Angelman syndrome^(2, 10, 11)^. These findings validate scalp EEG as a noninvasive and translationally relevant tool for assessing neural function in this large-animal model. The ability to capture human-like electrophysiological signatures in pigs provides a critical bridge between rodent and clinical studies and establishes a robust foundation for future therapeutic testing. Together, these results highlight the promise of the pig model for advancing preclinical research in Angelman syndrome and accelerating translation of potential treatments to clinical practice.

## Acknowledgements

We thank Ozair Habib, Morgan Matt, Thomas Jepp, Daniela Ramos, Aubrey Rinderknecht, Katie Bumgardner, and Milton Mosk for their contributions to this project. We also acknowledge the support of the staff in the Comparative Medicine Program and the Texas Institute for Preclinical Studies at Texas A&M University.

We acknowledge the use of ChatGPT (OpenAI) for assistance with proofreading and language refinement. The authors confirm that the model did not generate or influence any data, analyses, or scientific content and take full responsibility for the accuracy and integrity of the final manuscript.

